# Parallel Tempering with Lasso for Model Reduction in Systems Biology

**DOI:** 10.1101/720631

**Authors:** Sanjana Gupta, Robin E.C. Lee, James R. Faeder

## Abstract

Systems Biology models reveal relationships between signaling inputs and observable molecular or cellular behaviors. The complexity of these models, however, often obscures key elements that regulate emergent properties. We use a Bayesian model reduction approach that combines Parallel Tempering with Lasso regularization to identify minimal subsets of reactions in a signaling network that are sufficient to reproduce experimentally observed data. The Bayesian approach finds distinct reduced models that fit data equivalently. A variant of this approach based on Group Lasso is applied to the NF-*κ*B signaling network to test the necessity of feedback loops for responses to pulsatile and continuous pathway stimulation. Taken together, our results demonstrate that Bayesian parameter estimation combined with regularization can isolate and reveal core motifs sufficient to explain data from complex signaling systems.

## Introduction

Cells use complex networks of proteins and other biomolecules to translate environmental cues into various cell fate decisions. Mathematical and computational models are increasingly used to analyze the nonlinear dynamics of these complex biochemical signaling systems [1–4]. As our knowledge of the biochemical processes in a cell increases, reaction network models of cell signaling have been growing more detailed [4–6]. Detailed models are a useful summary of knowledge about a system but they suffer from several drawbacks. First, the complexity may obscure simpler motifs that govern emergent cellular functions [7–9]. Second, the large number of parameters creates a high-dimensional search problem for parameter values where the model fits the data. To mitigate these problems, it is useful to reduce the number of reactions in a model, provided that the reduced model is still able to reproduce a given set of experimental observations. In this work we pose model reduction as a constrained Bayesian parameter estimation (BPE) problem to simultaneously calibrate and reduce models. Given a prior reaction network model, our method finds all possible minimal subsets of non-zero parameters that fit the data.

Previous studies have addressed model reduction for biochemical systems [10]. Some examples include reductions by topological modifications to resolve non-identifiability in models [11, 12] and reductions by timescale partitioning [13, 14]. Non-identifiability arises when multiple unique parameterizations of a model give the same model output. Quaiser et al. [11] and Raue et al. [12] developed methods to find non-identifiable parameters and used this analysis to resolve non-identifiability by model simplifications such as lumping or removal of reactions. The simplification step, however, is not automated and requires a skilled modeler. Timescale partitioning methods use timescale separations in the reaction kinetics to apply model reduction based on quasi-steady-state and related approximations [10, 14]. Both of these methods generate reduced models but do not carry out parameter estimation to fit experimental data. Another approach to model reduction that includes parameter estimation [15] uses mixed-integer nonlinear optimization to combine parameter estimation with model reduction by reaction elimination, a technique common in the field of chemical engineering [16, 17]. Drawbacks of this approach are that it requires an additional binary parameter for every reaction in the model and that the genetic algorithms used for the optimization only provide point estimates of the parameters.

Here, we develop reaction elimination in a Bayesian framework that combines parameter estimation and model reduction without requiring additional parameters. BPE has been shown to be useful to characterize high-dimensional, rugged, multimodal parameter landscapes common to systems biology models [1, 18–21], but it suffers from the drawback that the Markov Chain Monte Carlo (MCMC) methods commonly used to sample model parameter space are often slow to converge and do not scale well with the number of model parameters. We recently showed that Parallel Tempering (PT), a physics-based method for accelerating MCMC [22], outperforms conventional MCMC for systems biology models with up to dozens of parameters [18]. Here, we apply Lasso (also known as L1 regularization), a penalty on the absolute values of the parameters being optimized, to carry out model reduction. In the fields of statistics and machine learning, Lasso is widely used for variable selection to identify a parsimonious model – a minimal subset of variables required to explain the data [23]. In the context of biology, Lasso has been widely applied to gene expression and genomic data typically in combination with standard regression techniques [24–28] and less commonly in Bayesian frameworks [29, 30]. In the mechanistic modeling context, Lasso regression has been used to predict cell type specific parameters in ODE reaction network models [31], but to our knowledge it has not been implemented to reduce such models.

Our method, PTLasso, combines PT with Lasso regularization to simultaneously calibrate and reduce models. The core idea is that every reaction in the model is governed by a rate constant parameter that, when estimated as zero, removes the reaction from the model simulation. Since the approach is Bayesian, PTLasso extracts all possible minimal subsets of reactions, which provides alternate mechanisms to explain the data. Using both synthetic and real biological data, we demonstrate that PTLasso is an effective approach for model reduction. We also apply PT coupled with Group Lasso [32] in a larger model of NF-*κ*B signaling to select over reaction-network modules instead of individual reactions. Group Lasso can test mechanistic hypotheses about the necessity of signaling modules, such as feedback loops, to explain data from particular experimental conditions. Overall, our results demonstrate that BPE combined with regularization is a powerful approach to dissect complex systems biology models and identify core reactions that govern cell behavior.

The remainder of this paper is organized as follows. In Methods we provide an overview of the PTLasso approach with in-depth descriptions of PT, Lasso regularization, Group Lasso and the setup of computational experiments. In Results we demonstrate PTLasso on synthetic examples of increasing complexity followed by an application of the Group Lasso approach to address mechanistic questions in NF-*κ*B signaling. Finally, in Discussion we highlight advances as well as limitations of the method and present the implications of this study for the broader context of biological modeling and analysis.

## Methods

In this work we use Bayesian parameter estimation (BPE) for model reduction. Following [18], BPE methods aim to estimate the probability distribution for the model parameters conditioned on the data. The probability of observing the parameter vector, 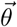, given the data, *Y*, is given by Bayes’ rule

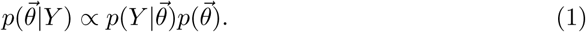

Here, 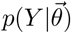 is the conditional probability of *Y* given 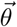, and is described by a *likelihood model*. For the ODE models in this study, we assumed Gaussian experimental measurement error, in which case the likelihood of a parameter vector, 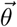, is given by

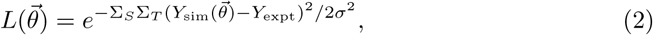

where *S* is a list of the observed species, *T* is a list of the time points at which observations are made, *σ* is the standard deviation of the likelihood model, 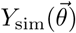 is the model output for parameter vector, 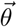, and *Y*_expt_ is the corresponding experimental data. 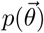 is the independent probability of 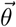, often referred to as the *prior distribution*, which represents our prior beliefs about the model parameters. It can be used to restrict parameters to a range of values or even to limit the number of nonzero parameters, as discussed further below.

### MCMC sampling

MCMC methods sample from the posterior distribution, *p*(*θ*|*Y*), by constructing a Markov chain with *p*(*θ*|*Y*) as its stationary distribution. Following the notation of Metropolis *et al*. [33], we define the energy of a parameter vector 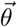 as

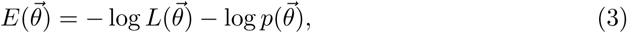

where *L* and *p* are the likelihood and prior distribution functions defined above. In this section we will briefly describe the Metropolis-Hastings and Parallel Tempering algorithms for MCMC sampling.

### Metropolis-Hastings algorithm

The Metropolis-Hastings (MH) algorithm is a commonly-used MCMC algorithm for BPE [34]. At each step, *n*, the method uses a proposal function to generate a new parameter vector, 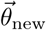, given the current parameter vector,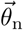. A common choice of proposal function is a normal distribution centered at 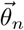:

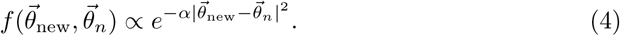

For any *f* that is symmetric with respect to 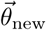 and 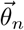, the move 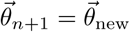 is accepted with probability min(1, *e*^−Δ*E*^), wheren 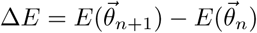. If the move is not accepted *θ*_*n*+1_ is set to *θ*_*n*_

### Parallel Tempering

In PT (also referred to as replica exchange Monte Carlo [22]), several Markov chains are constructed in parallel, each with a different temperature parameter, *β*, which scales the acceptance probability from the MH algorithm, which is now given by min(1, *e*^−*β*Δ*E*^) A Markov chain with *β* = 1 samples the true energy landscape as in MH. Higher temperature chains have *β <* 1 and accept unfavorable moves with a higher probability, sampling parameter space broadly. Tempering refers to periodic attempts to swap parameter configurations between high and low temperature chains. These moves allow the low temperature chain to escape from local minima and improve both convergence and sampling efficiency [22]. Following [18], the PT algorithm is as follows:

1. For each of *N* swap attempts (called “swaps” for short)
  a. For each of *N*_*c*_ chains (these can be run in parallel)
    i. Run *N*_MCMC_ MH steps
    ii. Record the values of the parameters and energy on the final step.
  b. For each consecutive pair in the set of chains in decreasing order of temperature, accept swaps with probability min(1, *e*^Δ*β*Δ*E*^), where Δ*E* and Δ*β* are the differences in the energy and *β* of the chains, respectively.

Adapting the step size and the temperature parameter can further increase the efficiency of sampling [22], but varying parameters during the construction of the chain violates the assumption of a symmetric proposal function (also referred to as “detailed balance”). It is therefore advisable to do this during a “burn-in” phase prior to sampling.

### Regularization with Lasso

Lasso regularization penalizes the L1-norm (sum of absolute values) of the parameter vector, which biases all model parameters towards a value of zero [23]. In a Bayesian framework, the Lasso penalty is equivalent to assuming a Laplace prior on each parameter *θ*_*i*_ given by

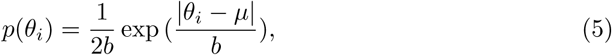

where *b* is the width and *µ* is the mean, which is set to zero for variable selection in linear parameter space. The energy function is then

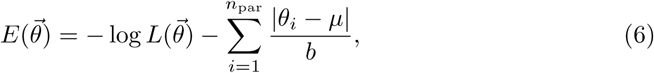

where *n*_par_ is the number of model parameters. Since PT only uses energy differences, the normalization constant 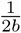 in Eq. 5 does not appear in Eq. 6. *b* is inversely proportional to the regularization strength.

For efficiency we usually perform parameter estimation in log parameter space, so instead of regularizing by setting *µ* to zero, we set it to a large negative value, such that the parameter value is small enough that it does not affect the dynamics of the model variables on the timescale of the simulation.

### Regularization with Group Lasso

To account for modularity in complex signaling networks [35], we use Group Lasso, previously described for regression problems [32], to perform selection at the level of reaction modules instead of individual reactions. All reactions in a module share a common penalty parameter that is multiplied with a reaction-specific parameter to get the full reaction rate constant.

For every reaction *i* in module *m*, the reaction rate constant is given by

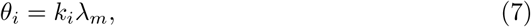

where *λ*_*m*_ is the penalty parameter for module *m* and *k*_*i*_ is a reaction-specific parameter. Defining 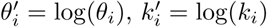, and 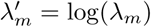, we have

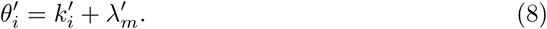

The energy function is then

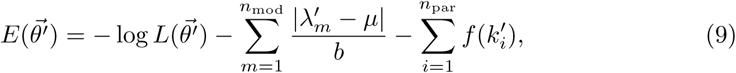

Where

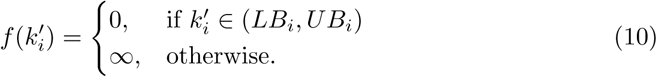

Here, *n*_mod_ is the number of modules and *LB*_*i*_ and *UB*_*i*_ are parameters that restrict the reaction-specific parameters. *UB*_*i*_ is chosen such that when 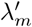 is within the Laplace prior boundaries, i.e 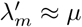, the maximum value of 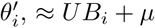, is small enough that it does not affect the dynamics of the model variables on the timescale of the simulation. For the application to NF-*κ*B signaling we chose *µ* = −25, *LB*_*i*_ = −5 and *UB*_*i*_ = 10 for all *i*.

### Synthetic data sets used in model calibration

For the two examples presented in Results that used synthetic data, we generated the sets labeled “true data” by simulating the model with a single set of parameter values and sampling with a fine time resolution. We then generated 10 noisy replicates of this data at a coarser set of time points by adding Gaussian noise with mean of zero and variance of either 10% or 30% of the true value at each point. The mean and variance of the replicates then defined the “observed data” used for fitting.

### Constraining the model

We use two kinds of constraints in fitting, soft constraints and hard constraints. Soft constraints can be violated, but are associated with a finite penalty [36]. For example, the energy function penalizes parameter vectors for producing model outputs that deviate from the data. Hard constraints, on the other hand, cannot be violated because they are associated with an infinite penalty. We used hard constraints in the NF-*κ*B signaling model to enforce certain known properties of the NF-*κ*B system, such as that the exit rate of NF-*κ*B -I*κ*B complex from the nucleus is greater than that of free NF-*κ*B [37]. A full list of constraints applied to the NF-*κ*B signaling model is listed in Table S4.

### MCMC chain initialization

All MCMC chains must be initialized with a starting parameter vector. For simple examples, such as the pulse-generator motif and linear dose-response models, chains were initialized by randomly sampling from the prior until a parameter vector with energy below a threshold is found. For more complex examples, to avoid long burn-in periods when starting from unfavorable start points, parameter vectors obtained from short PT runs were used to initialize longer PT chains. For example, parameter sets obtained for one NF-*κ*B trajectory could be used as a start point for fitting a different NF-*κ*B trajectory, or a PTLasso chain with a small value of *b* (more constrained) could be initialized from a parameter set obtained from PTLasso with a large value of *b* (less constrained). The exact procedures used to generate the starting configurations used in all computational experiments are provided in the Supplemental Code that will be provided on GitHub upon publication of this manuscript.

### Convergence testing

To check for convergence, PT or PTLasso was run twice for each computational experiment, and the two parameter chains were used to calculate the Potential Scale Reduction Factor (PSRF) for each model parameter. The PSRF compares intra-chain and inter-chain variances for model parameters and serves as a measure of convergence [38]. In keeping with the literature, we consider a PSRF less than 1.2 [19, 20, 38] as indicating convergence. We also calculated the stricter Multivariate PSRF (MPSRF), which extends PSRF by checking for convergence of parameter covariation. Third-party MATLAB libraries used for the MPSRF calculations are available at https://research.cs.aalto.fi/pml/software/mcmcdiag/.

For models with a large number of parameters, such as the 26-parameter NF-*κ*B signaling model, the number of PT samples needed for convergence was large and time consuming to obtain in a single run. Instead of running two long PT chains each of length *N*, we picked two favorable initial conditions and from each ran a set of *M* PT chains of length *N/M* in parallel to reduce wall clock time. We calculated the univariate PSRF of the *M* energy chains within each group, and if PSRF was less than 1.2, we assumed that the chains were sampling the same energy basin and combined them (Table S5). This gave us two groups of *N* PT samples that we used to calculate parameter convergence.

PSRF and MPSRF valuess for each computational experiment are shown in Tables S1 and S2 respectively. We also show in Table S3 that the acceptance rates for most chains are close to the optimal value of 0.234 [39].

### Hyperparameter selection

The hyperparameters associated with PTLasso are *µ* and *b*, the mean and width of the Laplace prior on each parameter that is being regularized. For simplicity, we keep these the same for all model parameters, although they could in principle vary, which would lead to a more difficult inference problem. To select the hyperparameters, we varied *b* and used the “elbow” in the negative log likelihood vs. *b* plot to find the smallest value of *b* (maximum regularization strength) that does not substantially increase the negative log likelihood of the fit [18, 40]. We also checked that the results were insensitive to small variations in *µ* (Figs. S1–S3).

For more computationally expensive models, we used hyperparameter estimates close to those obtained from the smaller synthetic models and compared the average log likelihoods of the fits from PT and PTLasso. For all of the examples shown, we found that the fit with PTLasso is at least as good as the fit with PT (Figs. 4, S1–S3).

### Software

All results reported in this work were obtained using `ptempest` [18], which is a MATLAB package for parameter estimation that implements PT with support for regularization. The source code is available at http://github.com/RuleWorld/ptempest.

## Results

### Reduced motifs can be inferred from dense reaction-networks in the absence of a prior architecture

To demonstrate that PTLasso can recover a minimal model architecture without prior knowledge of the reaction network, we used synthetic time-course data to infer a pulse-generator motif from a fully connected 3-node network of unimolecular reactions. The motif A → B → C (Fig. 1A, left), modeled as a system of ODE’s, was used to generate a time course for species B after initializing the system with 100 molecules of species A at time *t* = 0 (red curves in Fig. 1B labeled ‘true data’). To simulate the effects of experimental noise and cell-to-cell variability, Gaussian noise was added to generate ten noisy trajectories that were sampled at eight time points (Fig. S1A). The mean and standard deviation of these synthetic trajectories formed the ‘observed data’ (black points and error bars in Fig. 1B) used for subsequent parameter estimation and model reduction.

**Fig 1.**
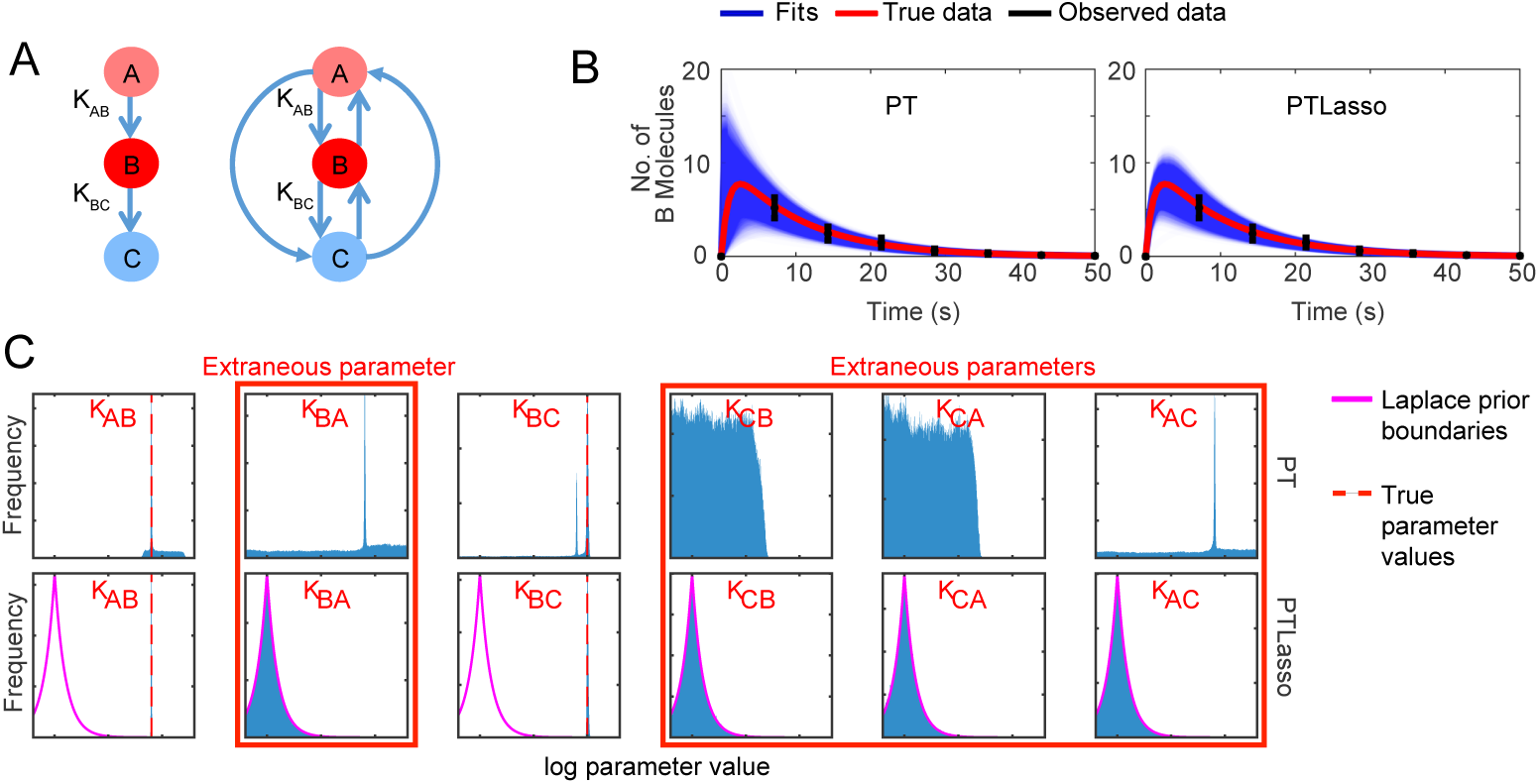
Model reduction using Lasso with a fully connected 3-node graph. **A)** Motif used to generate the observed data (left) and reaction network diagram of the fully connected 3-node network used as the starting point for PTLasso (right). The initial concentration of A (light red) is 100 molecules. The initial concentrations of B and C are 0. The concentration of B (red) is observed at multiple time points, but the concentration of C (blue) is not observed. Each reaction has an associated rate constant parameter. *K*_*AB*_ = 0.1 and *K*_*BC*_ = 1 are the rate constant values used to generate the observed data. **B)** Fits of the model to the data with PT (left) and PTLasso (right). Blue lines show ensemble fits (4e3 samples, 100 time points per sample), red line shows the true data (100 time points), and the black error bars show the mean ± standard deviation of the observed data (8 time points). **C)** Frequency histograms showing probability distributions of the parameters (from 4e5 PT samples) for fits with PT (top row) and PTLasso (bottom row). The y-axis for each panel is scaled from 0 to the max value of the distribution to emphasize differences in the shapes of the distribution. The pink lines show the Laplace prior boundaries, the dashed red lines (panels for *K*_*AB*_ and *K*_*BC*_) show the parameter values of the known model used to generate the true data, and the x-axis range, [-12,3], represents the sampling range. Parameter distributions confined within the Laplace prior boundaries are extraneous.

PT and PTLasso were then used to fit this data using the fully-connected 3-node network comprised of six reactions (Fig. 1A, right). Time courses from PT and PTLasso (Fig. 1B) both fit the observed data (Fig. S1C), but the PTLasso curves fit the true data better at times before the first observed data point. PT finds parameter probability distributions (Fig. 1C, top row) that exhibit sharp peaks near the exact values of the two nonzero parameters, K_AB_ and K_BC_, but finds significant probability for other values of these parameters and non-zero values for the other rate constants in the complete network that should have zero value (labeled ‘extraneous’). By contrast, PTLasso (Fig. 1C, bottom row) recovers tight distributions on the values of the two nonzero parameters that lie well outside the Laplace prior, while the probability distributions for the extraneous parameters all conform tightly to the prior distribution, indicating that the corresponding reactions can be removed from the network. Taken together, these results demonstrate that PTLasso can recover network architecture and parameter values that are not inferred by PT alone.

To determine if the method scales to larger networks, we applied PT and PTLasso to a fully connected 5-node network (Fig. 2A, Fig. S2A). As with the 3-node example, PTLasso fits for a complete 5-node network are more similar to the true data than fits with PT alone (Fig 2B). Similarly, rate constant parameter distributions with PT are all broad (Fig. 2C), whereas the extraneous parameters for the PTLasso fits were within the Laplace prior (Fig 2D). In addition to a correct reaction rate constant distribution for K_AB_, PTLasso recovered bimodal distributions for and K_BC_, K_BD_, and K_BE_, suggesting that the essentiality of each of the reactions B → C, B → D and B → E depends on which of the other two are included. This is because the model A → B → C is indistinguishable from A → B → D and A → B → E without more information about the system. Even though the marginal posterior distributions show all three parameters playing a role, parameter covariation (Fig. S2D, right) reveals that only one of the reactions B → C, B → D, B → E is simultaneously active and rate constant distributions for the other two are centered at 10^−10^ (proxy for 0 when sampling in log-scale). The same covariation plot obtained without Lasso does not show any meaningful structure (Fig. S2D, left). Taken together, these results show that PTLasso correctly identifies network parameters and suggests that A → B → C, A → B → D, and A → B → E are equivalent parsimonious models.

**Fig 2.**
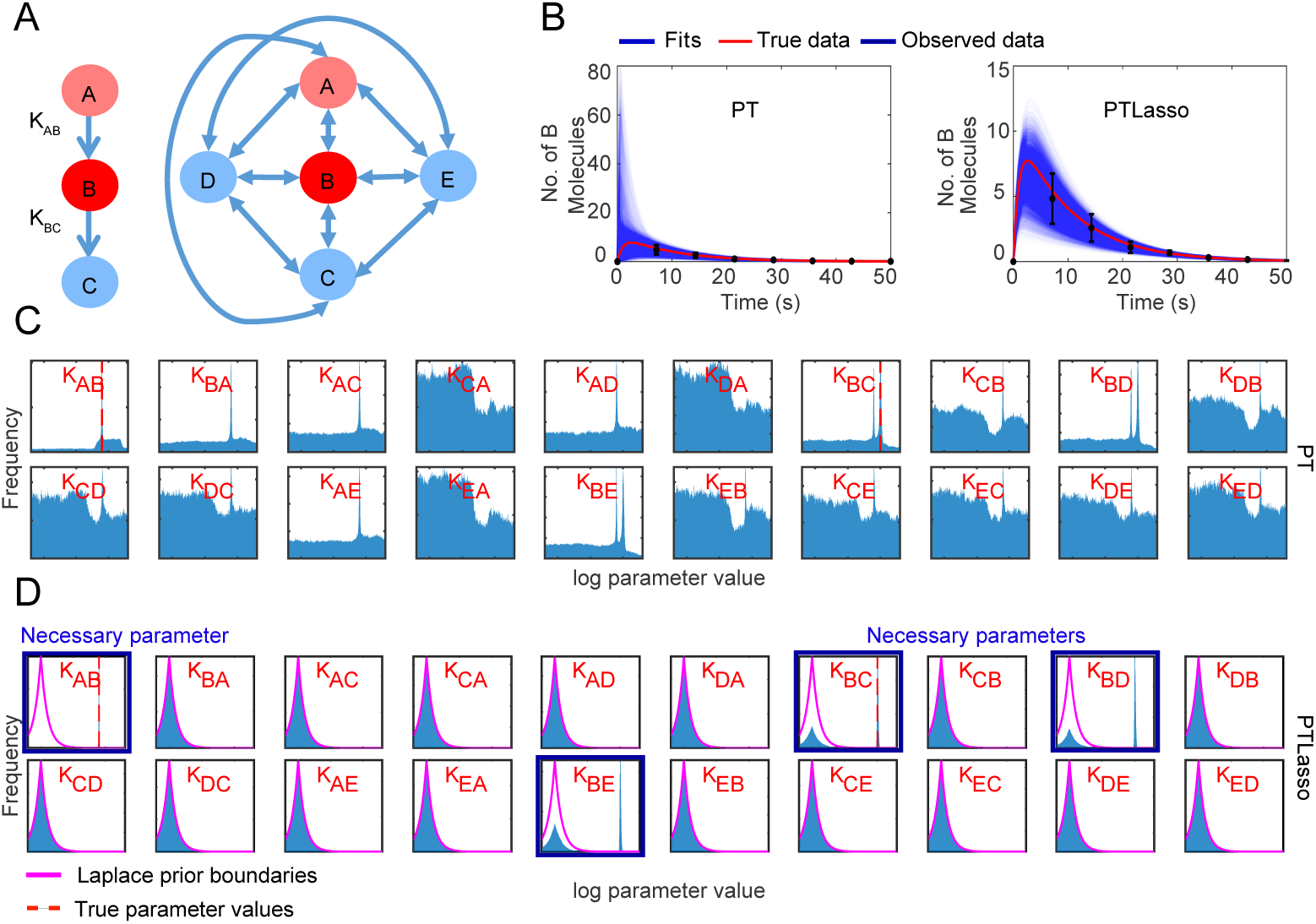
Model reduction using Lasso with a fully connected 5-node graph. **A)** Motif used to generate the observed data (left) and reaction network diagram of the fully connected 5-node network used as the starting point for PTLasso (right). The initial concentration of A (light red) is 100 molecules. The initial concentrations of B,C and D are 0. The concentration of B (red) is observed at multiple time points, but the concentration of C, D and E (blue) are not observed. *K*_*AB*_ = 0.1 and *K*_*BC*_ = 1 are the rate constant values used to generate the observed data. **B)** Fit of the model to the data with PT (left) and PTLasso (right). Blue lines show ensemble fits (7e3 samples, 100 time points per sample), red line shows the true data (100 time points), and the black error bars show the mean ± standard deviation of the observed data (8 time points). **C)** Frequency histograms showing probability distributions of the parameters (from 7e5 PT samples) for fits with PT and **D)** PTLasso. The y-axis for each panel is scaled from 0 to the max value of the distribution to emphasize differences in the shapes of the distribution. The pink lines show the Laplace prior boundaries, the dashed red lines (panels for *K*_*AB*_ and *K*_*BC*_) show the parameter values of the known model used to generate the true data, and the x-axis range, [-12,3], represents the sampling range. Parameter distributions that deviate from the prior are necessary.

Overall, our results show that PTLasso is a global approach that can extract correct parameter estimates and architectures of all equivalent reduced models that fit the data from fully connected networks of varying sizes. This is especially useful in the context of complex cell signaling systems that often have redundant elements in which case the method can be used to identify alternate signaling mechanisms that fit the data.

### Motifs with specific dose-response relationships can be inferred from a prior network

In the previous section we assumed no prior knowledge of a reaction-network and fitted a simple model output. To demonstrate the extraction of motifs with more complex behaviors in the more likely scenario where there is some prior network of hypothesized molecular interactions, we used PTLasso to extract parts of the prior network required to produce specific dose-response relationships.

Tyson et al. [9] previously described two simple biochemical models that individually produce linear or perfectly adapting dose-response relationships. We constructed a prior network of a signal, S, response, R, and intermediate, X, by combining the linear and adaptive dose-response models into a single shared architecture (Fig. 3A). We show that PTLasso correctly identifies the linear and adaptive submodels when the combined model is fit to different simulated data. To begin with, the linear dose-response submodel was used to generate synthetic time courses for R in response to 4 increasing levels of S (S=1,2,3,4). As earlier, Gaussian noise was added to each trajectory to simulate experimental noise and cell-to-cell variability, and the mean and standard deviation for each time course was calculated at 4 distinct time points (including t=0), creating 16 data points that constitute the observed data. When fit to the observed data, PT produced fitted time courses for R that go through the observed data points but have fast time scale deviations from the true data time courses (Fig. 3B, left). PTLasso again produced fits closer to the true data (Fig. 3B, right). Parameter distributions inferred with PTLasso show that the 2-parameter model reduced from the combined model correctly recovers the architecture and parameterization of the linear-dose response submodel used to generate the synthetic time courses (Fig 3C,E).

**Fig 3.**
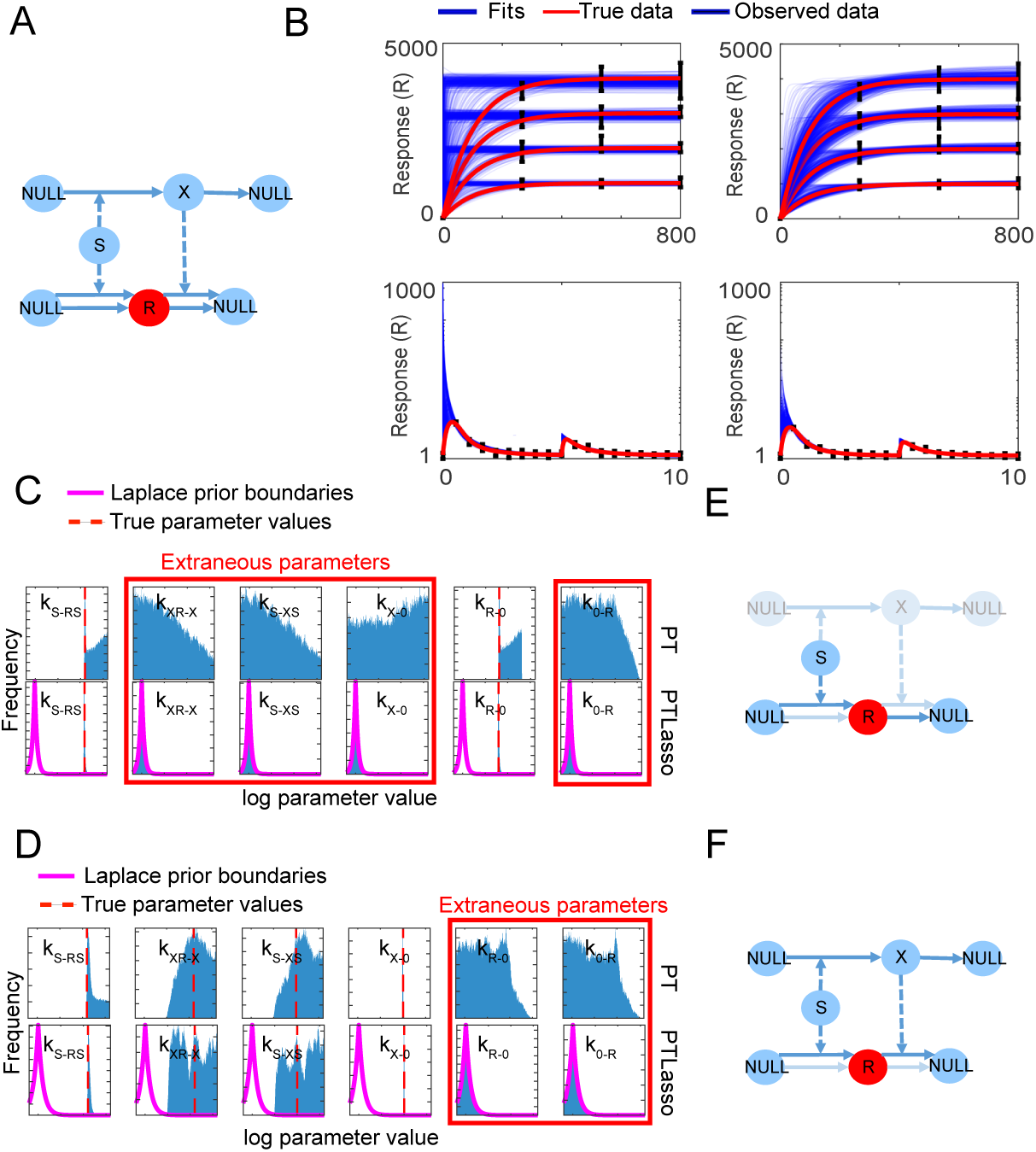
Motif inference from a prior network constrained with dose-response data. **A)** Reaction network diagram of the prior network. The value of the signal S is known, response R (red) is observed at multiple time points, but intermediate X (blue) is not observed. Solid lines show species conversions and dashed lines show influences. *k*_*s*−*rs*_ = 10, *k*_*r*−0_ = 0.01 are the rate constant values used to generate observed data for the linear dose-response model. *k*_*s*−*rs*_ = 10, *k*_*xr*−*x*_ = 10, *k*_*s*−*xs*_ = 1, *k*_*r*−0_ = 1 are the rate constant values used to generate observed data for the perfectly adapting dose-response model **B)** Fit of the model to the linear dose-response data (top, y-axis is in linear scale) and perfectly adapting dose-response data (bottom, y-axis is in log scale) with PT (left) and PTLasso (right). Blue lines show ensemble fits (400 samples with 1000 time points for linear dose-response, 800 samples with 2000 time points for perfectly adapting dose response), red line shows the true data (1000 time points for linear dose-response, 2000 time points for perfectly adapting dose response), and the black error bars show the mean ± standard deviation of the observed data **C)** Frequency histograms showing probability distributions of the parameters for linear dose response fits (from 4*e*5 PT samples) and **D)** perfectly adapting dose-response (from 8*e*5 PT samples) with PT (top) and PTLasso (bottom). The pink lines show the Laplace prior boundaries, the dashed red lines show the parameter values of the known model used to generate the true data, and the x-axis range, [-12,6], represents the sampling range. Parameter distributions that deviate from the prior are necessary, while those that are confined within the Laplace prior boundaries are extraneous. **E)** Reduced model corresponding to linear dose-response and **F)** perfectly adapting dose-response highlighted in prior network. Faded nodes and arrows are extraneous and are removed from the model. Solid lines show species conversions and dashed lines show influences.

When the perfectly adaptive dose-response submodel was similarly used to generate observed data in response to two successive increasing signal concentrations, PTLasso reduced the combined system to a 4-parameter model (Fig 3D,F). In this case, parameters *k*_*s*−*xs*_ and *k*_*xr*−*x*_ in the reduced model are unidentifiable, however, PTLasso captures their linear correlation (Fig. S3A) even though these parameters are not individually constrained, providing a complete picture of the system. While signaling systems are complex and can involve large numbers of reactions, not every reaction is relevant for every function. Taken together our results demonstrate that distinct elements of a large reaction-network may be responsible for different complex behaviors and can be successfully isolated using PTLasso.

### A reduced model of NF-*κ*B signaling without A20 feedback explains single-cell NF-*κ*B responses to a short TNF pulse

Complex biological signaling networks are frequently modular [41, 42] with distinct motifs such as feedback loops that operate on separate time scales [35]. To account for the modular structure of signaling we extended our Lasso approach to Group Lasso [32], a technique that applies a module-specific Lasso penalty to all reactions within a particular module (see Methods). PT combined with Group Lasso finds minimal sets of reaction modules that explain experimental data. We used this method to test the requirement of A20 feedback to explain previously published single-cell NF-*κ*B responses to a short TNF pulse [43].

A prior model of NF-*κ*B signaling was created by combining simplified elements of models from [37] and [2]. The network was divided into three biologically motivated network modules (Fig. 4A). The I*κ*B and A20 modules describe negative feedback mediated by the inhibitor I*κ*B and negative regulator A20, respectively. The activation module includes all remaining reactions that describe the path from TNF binding to its cognate TNF-receptor (TNFR) to the eventual translocation of NF-*κ*B into the nucleus. The reaction rate constants within a module are constrained by a common Lasso penalty parameter (see Methods). If the penalty parameter for a module is estimated as 0, (here, 10^−25^ is used as a proxy for 0 when sampling in log scale), the entire module is removed from the simulation. To test which of the three modules are necessary to explain NF-*κ*B responses to a single TNF pulse, PTLasso was used to fit the model to three previously published, experimentally obtained, single-cell NF-*κ*B responses (Fig. 4B, Column 1) [43]. In addition to the NF-*κ*B data, other constraints were applied to make the system behave consistently with known biology. These constraints are listed in Table S4, and Fig. S4 demonstrates that PTLasso correctly followed the imposed parameter covariation.

**Fig 4.**
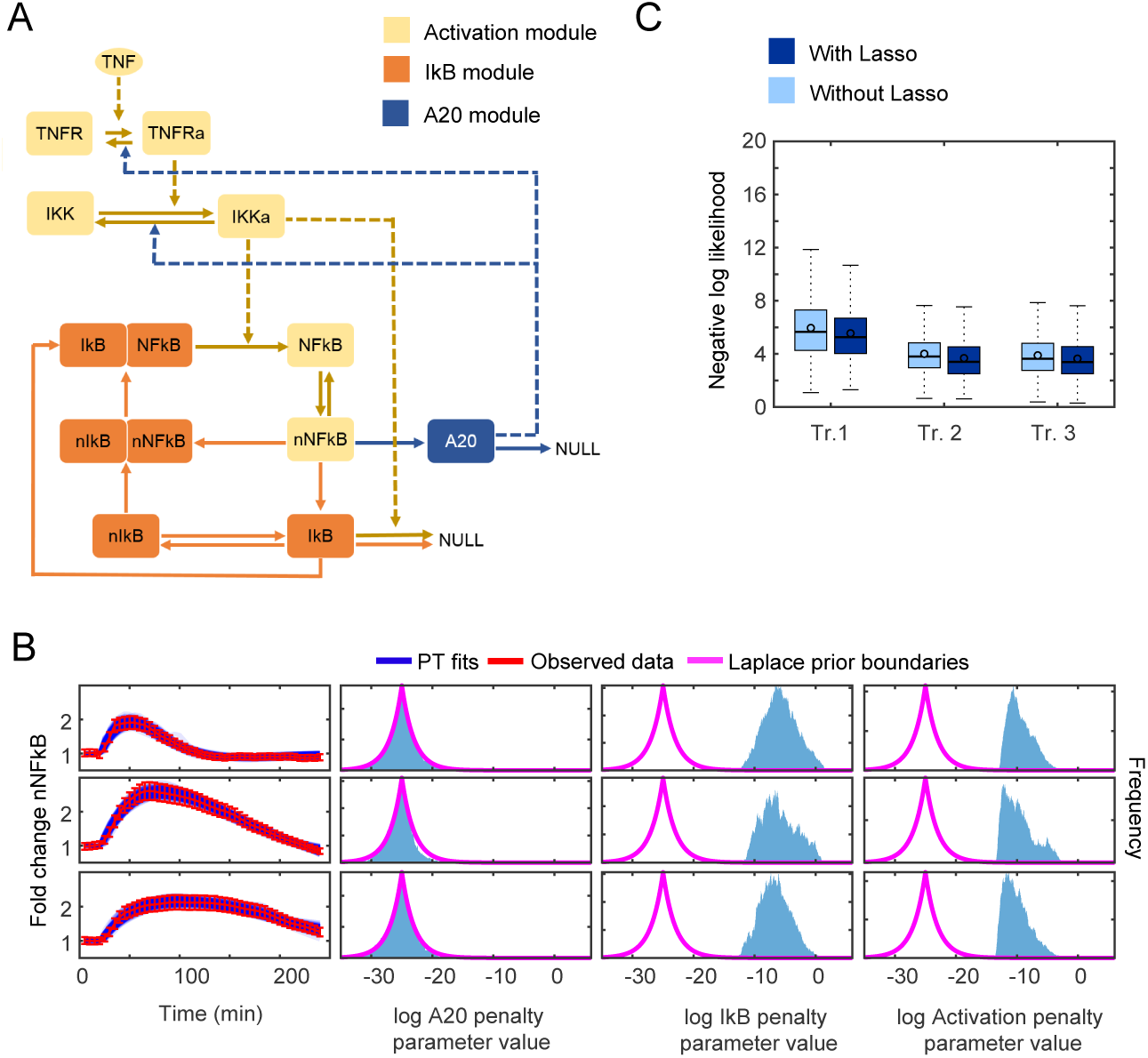
Model reduction using Group Lasso with a model of NF-*κ*B signaling. **A)** Reaction network diagram of a simplified model of TNF-NF-*κ*B signaling. The colors indicate the different modules. ‘a’ or ‘n’ refer to active and nuclear versions of the species respectively. Solid lines indicate transformations and dashed lines indicate influences. **B)** Ensemble fits of the model (in blue, 288 PTLasso samples, *µ*=-25,*b*=2) to a single-cell NF-*κ*B response to pulsatile TNF stimulation (in red). Error bars show the 10% standard deviation assumed for the likelihood function during fitting and represent measurement error (Column 1). Frequency histograms (from 5.64e6 samples) for the penalty parameter distributions corresponding to the different network modules (Columns 2-4). The pink lines show the Laplace prior boundaries, and the x-axis range, [-35,6], represents the sampling range. **C)** Box plots (5640 samples) comparing the log likelihood of the fits with PT and PTLasso (*µ*=-25,*b*=2) for the three trajectories (Tr. 1-3)

The probability distributions for the module penalty parameters (Fig. 4B, Columns 2-4) show that the A20 penalty parameter is confined within the prior boundaries while the others have deviated, suggesting that to fit these particular single-cell NF-*κ*B trajectories, the A20 module is dispensable, whereas the I*κ*B and activation modules are not. The A20 module might still be essential for other biology of the system, but the model does not require the A20 module to produce these single-cell NF-*κ*B responses under the given experimental condition and network constraints. The fits with PTLasso were as good as the fits with PT alone (Fig. 4B, Column 1), as is demonstrated by comparing the average log-likelihoods (Fig. 4C).

To test the requirement of A20 feedback under different experimental conditions and network constraints, we fit the model to a published single-cell NF-*κ*B response to continuous TNF stimulation [43]. A soft constraint that IKK responses are transient was added for consistency with published observations [44, 45]. For responses to a TNF pulse, IKK activity naturally adapts back to its baseline abundance without additional negative regulation (Fig. S6). In this case, all three module penalty parameters deviate from the prior (Fig. S7), indicating that the A20 mediated negative regulation of IKK is essential for responses to continuous TNF stimulation. Taken together, the results for the NF-*κ*B signaling model provide an example where PTLasso isolate reaction modules sufficient for responses to specific experimental conditions and time scales.

## Discussion

In this work we have demonstrated that PT combined with Lasso is an effective approach to learn reduced models from a prior model with a larger number of reactions. Even when starting from a complete graph without prior knowledge of the underlying signaling network, PTLasso correctly identified reduced model architectures and reaction rate constants. PTLasso also correctly isolated subnetworks that are necessary for distinct concentration and temporal dose-response relationships. In a model of NF-*κ*B signaling, PT with Group Lasso found that in the absence of other network constraints, A20 feedback was not required to explain single-cell responses to a short TNF pulse, but is required when TNF treatment was continuous. Model reduction using PTLasso can therefore highlight aspects of the reaction network that are important for specific experimental conditions and timescales and not others.

A potential application of model reduction arises in fitting a model to data from different cell types. Differences in responses to the same experimental condition might be explained by differences in parameters values [31], but comparing cell-type specific parameter distributions in high dimensional space may be difficult. Reducing the number of model parameters lowers the dimensionality of the space and makes this problem easier.

A limitation of PTLasso is the large number of samples required to reach convergence, which can lead to long execution times. For the simplest examples presented here, convergence happens on the order of hours on a standard workstation computer, but for the more complex signaling systems, convergence can take several days. Most of the execution time is dedicated to converging the joint parameter distribution. Currently PT and PTLasso are both run for fixed chain lengths followed by convergence testing at the end, often generating more samples than were required to pass convergence tests. Testing convergence on-the-fly and terminating PT chains when convergence is reached would prevent unnecessary sampling and reduce the overall execution time. Approaches such as APT-MCMC [46] and Hessian-guided MCMC [20] that account for the shape of the parameter landscape during sampling could also reduce the number of samples required for convergence.

Along with working to reduce the amount of sampling, we are also investigating algorithmic modifications to reduce the execution time of individual samples or PT swaps. Synchronous swapping in our current implementation of PT requires each chain to complete a fixed number of steps before attempting a swap. Because high temperature chains sample parameter space broadly and encounter regions where stiffness leads to long integration times, lower temperature chains often have to wait for the higher temperature chains to complete before swaps can be attempted. Asynchronous swapping [18] may therefore reduce execution times. Overall, there are still many opportunities for future PTLasso implementations to increase efficiency and applicability to larger systems biology models.

In this study we have presented a Bayesian framework that systematically dissects mechanistic ODE models of biochemical systems to identify minimal subsets of model reactions that are sufficient to explain experimental data. Technology now enables the building and simulating of highly detailed models that provide accurately reflect existing knowledge of a biochemical system. But detailed models may obscure our ability to identify underlying mechanisms. PTLasso serves as a bridge between these detailed models and simpler mechanistic explanations that can account for system behavior under specific conditions.

## Supporting information

**Fig S1.**
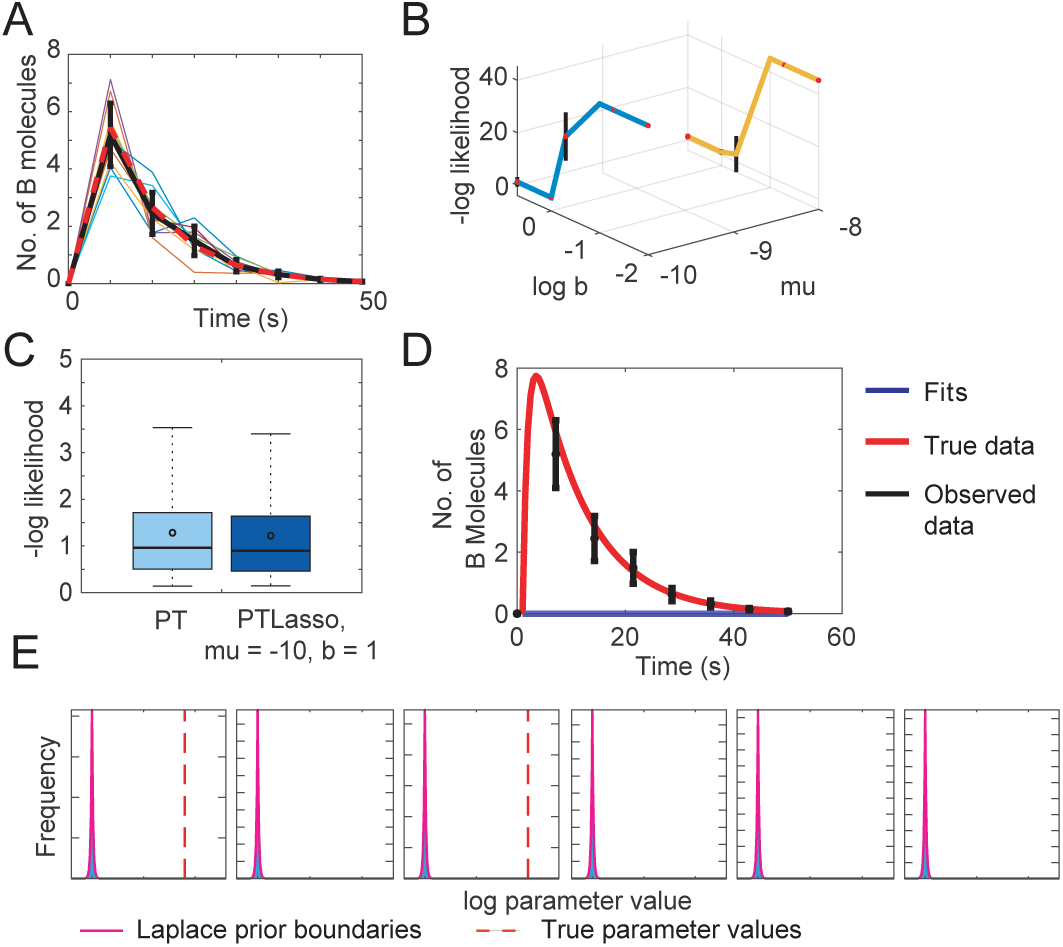
Hyperparameter tuning for PTLasso with a fully connected 3-node graph. **A)** Data generated for fitting. Red dashed lines show the model simulation at 8 time points with the known parameter values. Each colored line represents a noisy trajectory obtained by adding Gaussian noise to the true simulation. The black error bars show the mean and standard deviation of the 10 repeats, and is used for fitting. **B)** Hyperparameter tuning plot showing variation in the negative log likelihood distribution with *µ* and *b* (red points show the mean, and black lines show mean ± standard deviation for 4e3 samples). The hyperparameters selected (*µ* = −10, b = 1) provide the most regularization while not substantially increasing the negative log likelihood. **D)** Comparison of the log likelihood distributions (4000 samples) of the fits with PT and PTLasso (*µ* = −10, b = 1). Box plots are obtained using a third party MATLAB library, aboxplot*, with outliers not shown. **D)** Example of PTLasso fits (4e3 samples) where *b* is too small (*µ* = −10, b = 0.1) and the negative log likelihood of the fit is increased, and **E)** the corresponding parameter distributions (4e5 samples). Since the regularization strength was too high, none of the parameters deviated from the prior. *http://alex.bikfalvi.com/research/advanced_matlab_boxplot/

**Fig S2.**
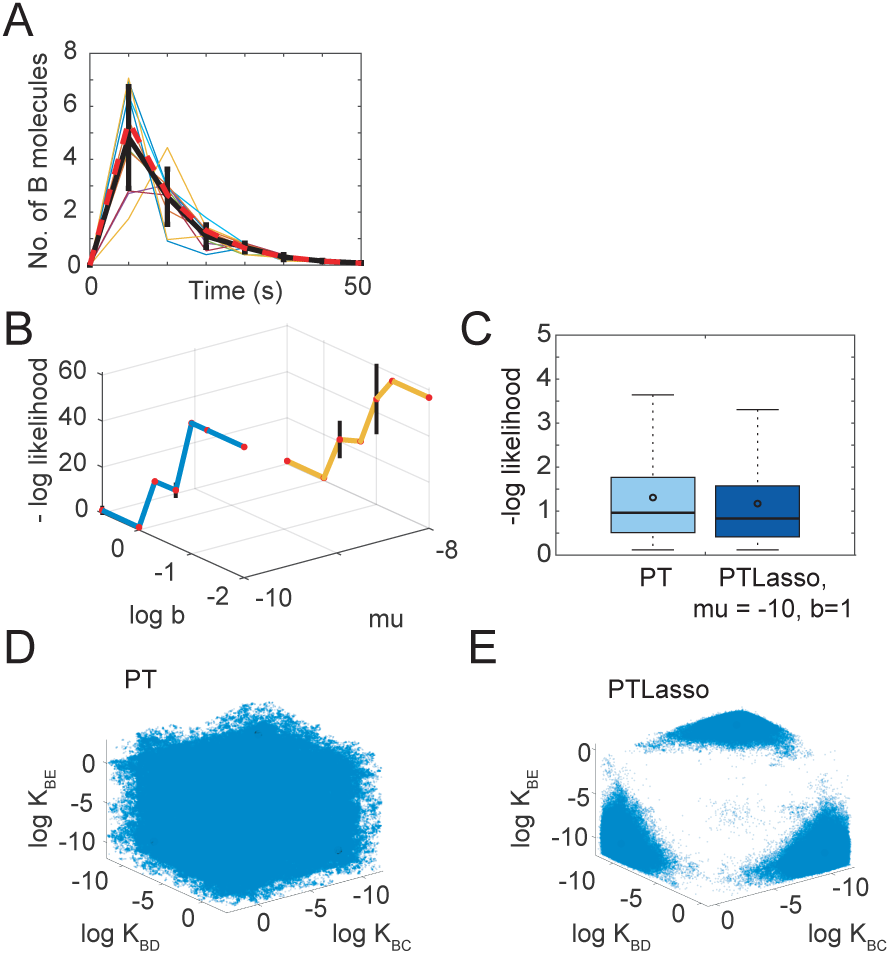
Hyperparameter tuning for PTLasso with a fully connected 5-node graph. **A)** Observed data generated for fitting. Red dashed lines show the model simulation at 8 time points with the known parameter values. Each colored line represents a noisy trajectory obtained by adding Gaussian noise to the true data. The black error bars show the mean and standard deviation of the 10 repeats, and is the observed data used for fitting.**B)** Hyperparameter tuning plot showing variation in the negative log likelihood distribution with *µ* and *b* (red points show the mean, and black lines show mean ± standard deviation for 7e3 samples). The hyperparameters selected (*µ* = −10, b = 1) provide the most regularization while not substantially increasing the negative log likelihood. **C)** Box plots comparing the log likelihood distribution (4000 samples) obtained with and without Lasso for the chosen values of hyperparameters. Box plots are obtained using a third party MATLAB library, aboxplot*, with outliers not shown. **D)**. Parameter co-variation (400000 samples from the distribution) of the three selected parameters with PT and **E)** with PTLasso. *http://alex.bikfalvi.com/research/advanced_matlab_boxplot/

**Fig S3.**
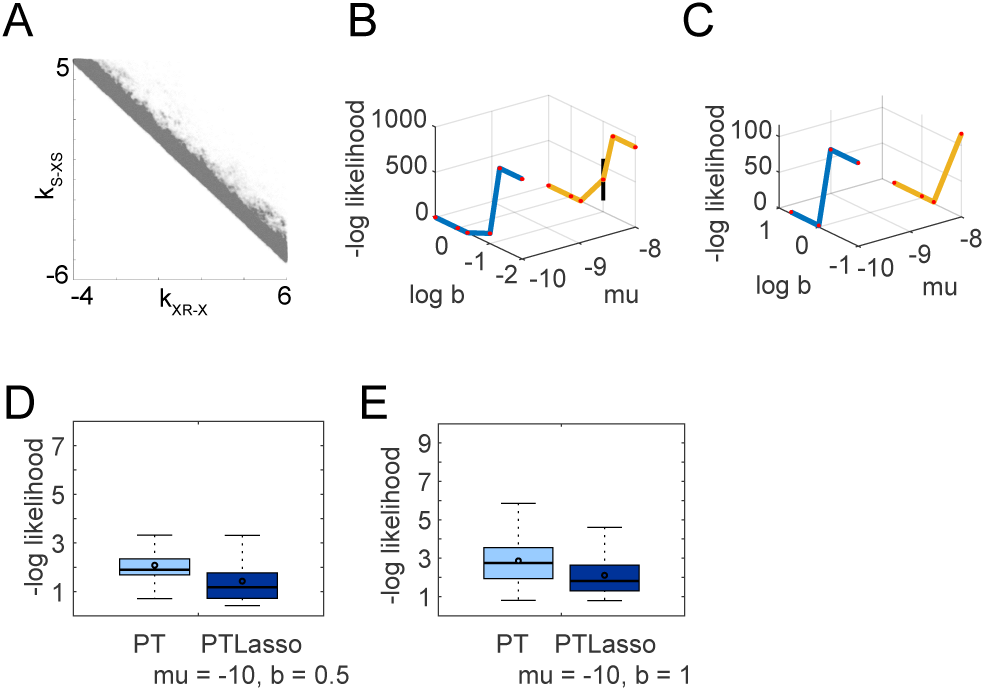
Hyperparameter tuning for PTLasso with dose-response motifs inferred from a prior network. **A)**. Linear correlation of non identifiable parameters in the reduced perfectly adapting model shown as a scatter plot (8e5 PTLasso samples). **B)** Hyperparameter tuning plot for the linear dose response model and **C)** the perfectly adapting dose response model. The hyperparameter tuning plot shows variation in the negative log likelihood distribution with *µ* and *b* (red points show the mean, and black lines show mean ± standard deviation, 4e2 samples for the linear dose response model and 8e2 samples for the perfectly adapting dose response model). The hyperparameters selected (*µ* = −10, b = 0.5 for linear dose-response and *µ* = −10, b = 1 for perfectly adapting dose-response) provide the most regularization while not substantially increasing the negative log likelihood. **D)** Box plots comparing the log likelihood distribution obtained with PT and PTLasso for the chosen values of hyperparameters for the linear dose response model (4e2 samples) and **E)** the perfectly adapting dose response model (8e2 samples). Box plots are obtained using a third party MATLAB library, aboxplot*, with outliers not shown. *http://alex.bikfalvi.com/research/advanced_matlab_boxplot/

**Fig S4.**
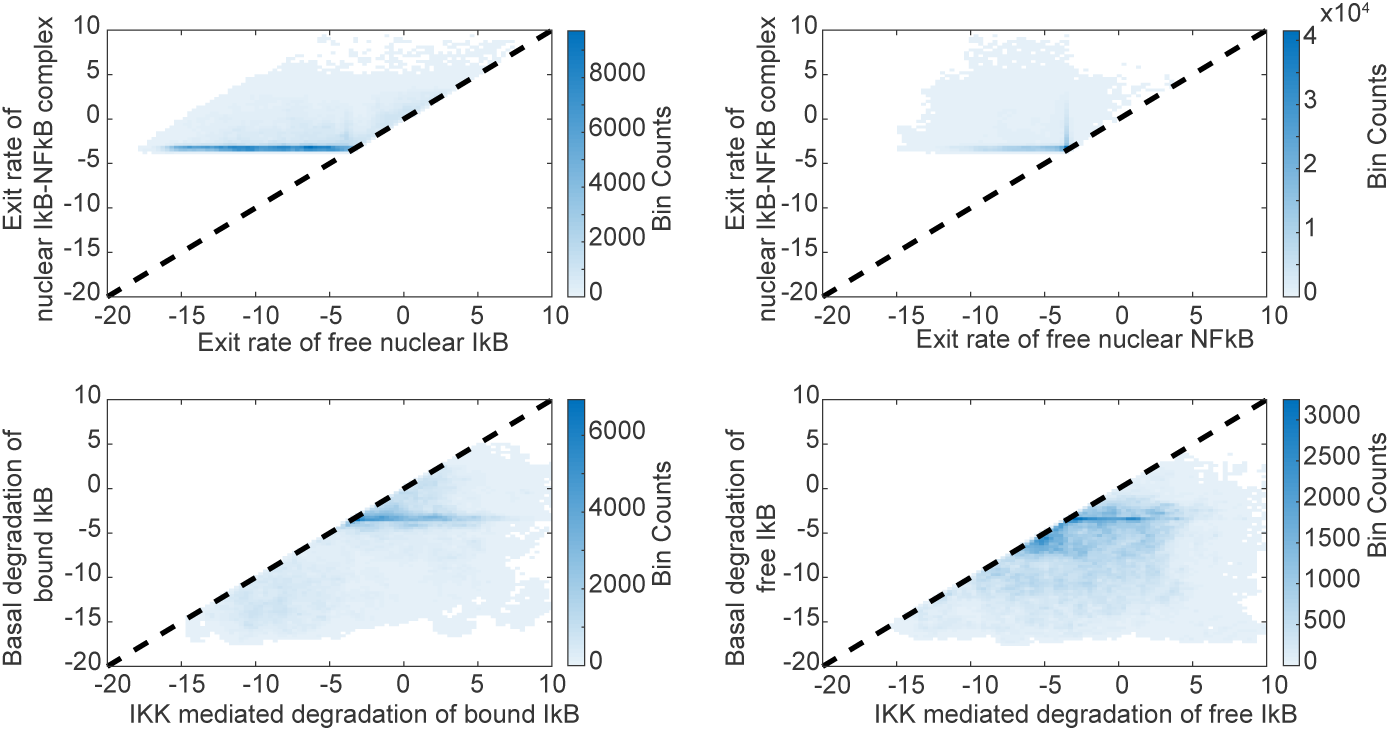
Constraints on parameter covariation in NF-*κ*B signaling. Binned scatter plots (MATLAB function binscatter with 9.4e5 samples from one of the PTLasso repeats) show the joint distributions for the pairs of parameters for which covariance was constrained during fitting.

**Fig S5.**
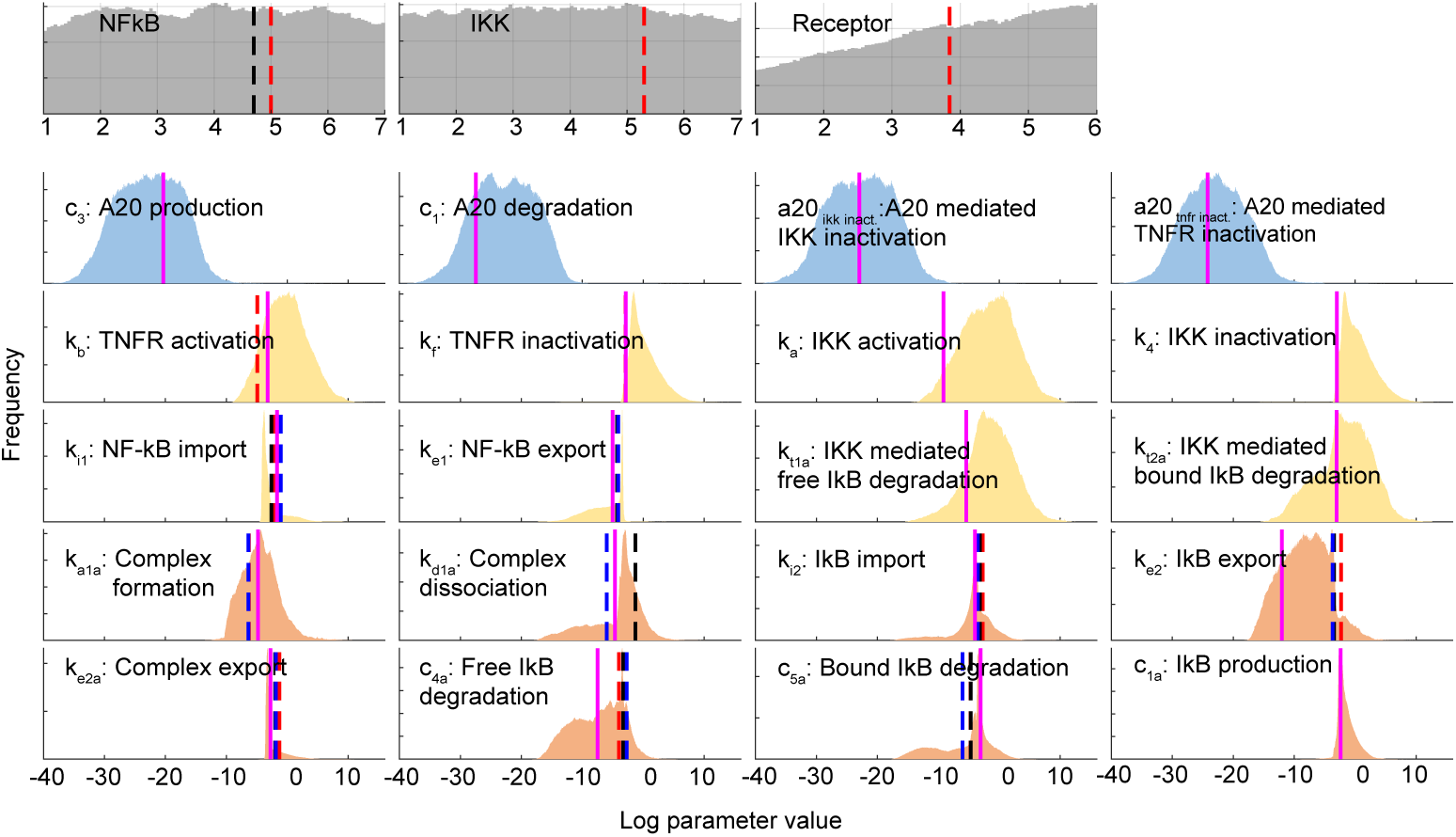
Marginal distributions of model parameters shown with the corresponding published values for a representative NF-*κ*B trajectory. Distributions of protein abundance parameters and rate constant parameters (from 5.64e6 samples) from the A20 module (blue), Activation module (yellow) and I*κ*B module (orange). All parameters are in logscale. Protein abundance parameters have uniform priors, and the x-axis range indicates the sampling range. Rate constant parameters are sums of the module penalty parameters and reaction-specific parameters. The pink line corresponds to a best-fit parameter set from one of the PTLasso repeats. The dashed lines correspond to published values of parameters – Pekalski et al. [2] (red), Lee et al. [37] (black), and Kearns et al. [47] (blue). A published parameter value for a model is only included if the corresponding reaction maintained the same structure in both the published and current models. For unit conversions we used the values mentioned in Lee et al., 1*µM* of NF-*κ*B = 50,000 molecules/cell and applied this to other species in the model. In the parameter labels ‘import’ refers to translocation from the cytoplasm into the nucleus and ‘export’ is the reverse. ‘Complex’ refers to the NF-*κ*B -I*κ*B complex.

**Fig S6.**
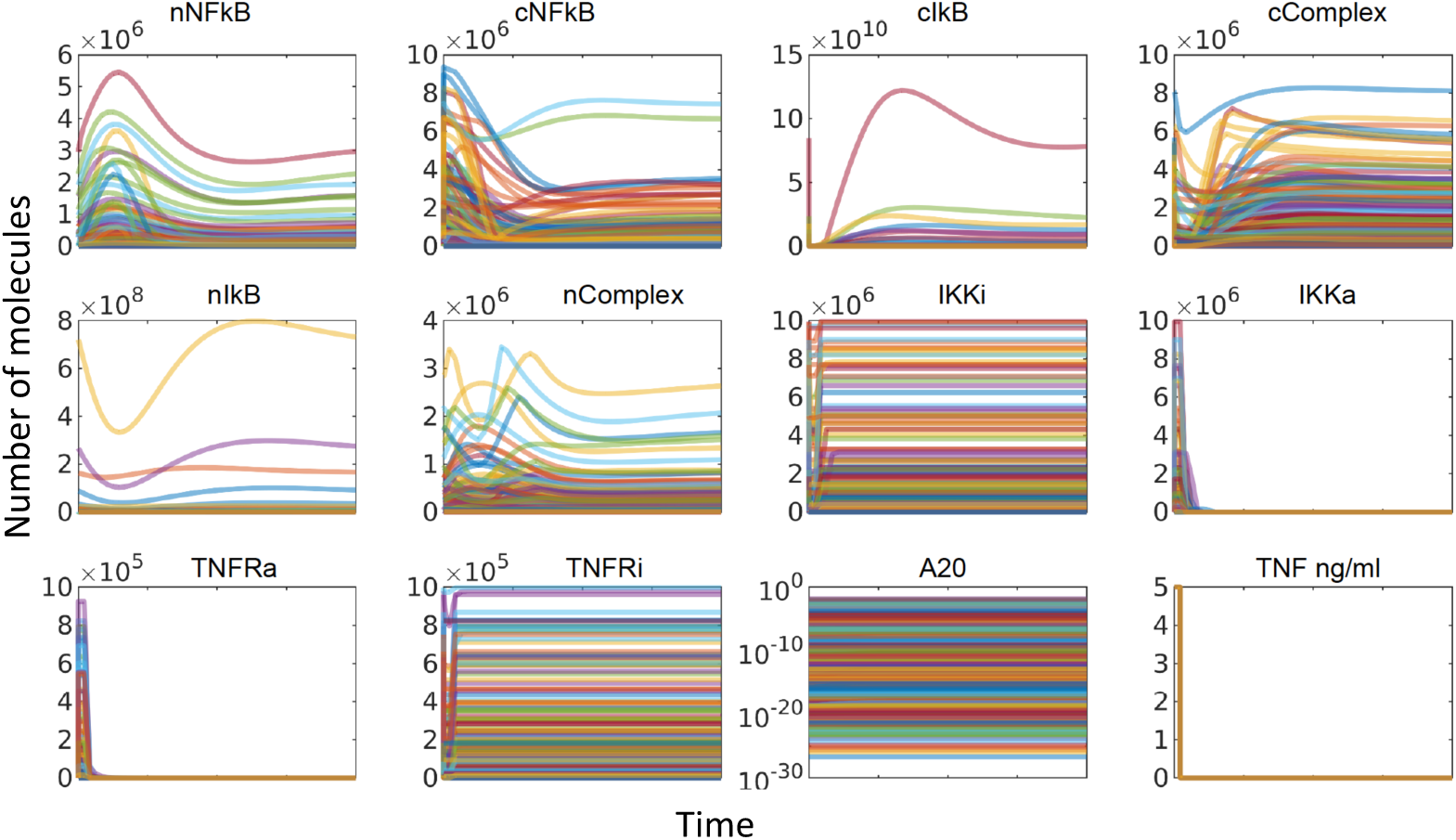
NF-*κ*B signaling model predictions. Model predictions for non-fitted variables for a representative NF-*κ*B trajectory (1881 samples from one of the PTLasso repeats). Time courses are shown for 240 minutes. Suffix ‘a’ and ‘i’ refer to active and inactive versions of a species respectively. Prefix ‘c’ and ‘n’ refer to cytoplasmic and nuclear versions of a species respectively. ‘Complex’ refers to the NF-*κ*B

**Fig S7.**
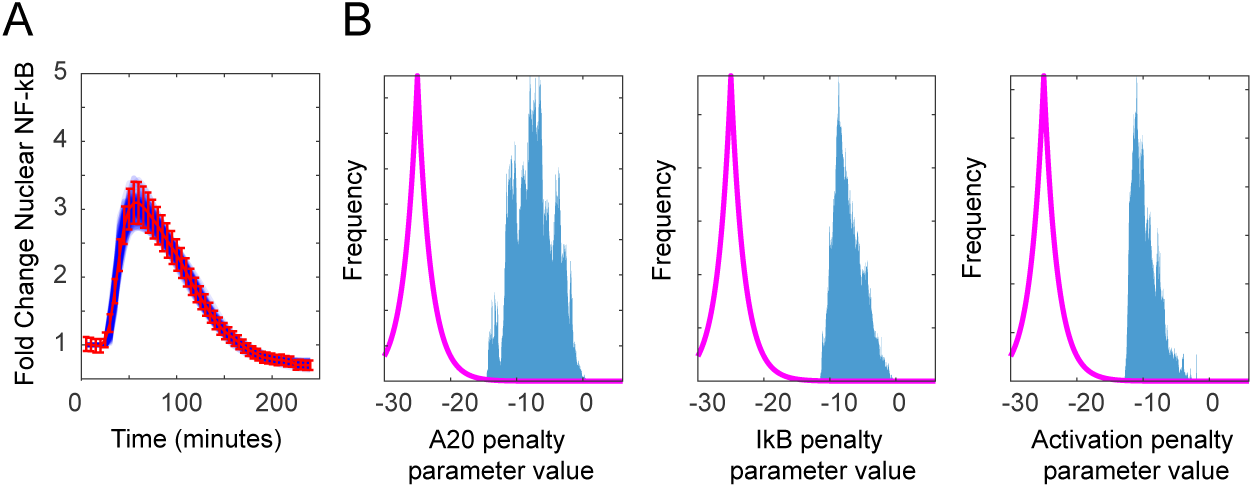
Group Lasso applied to NF-*κ*B signaling with continuous TNF stimulation. **A)** PTLasso fits of the model (in blue, 172 samples) to a single-cell NF-*κ*B response to TNF stimulation (in red). Errorbars show mean and standard deviation that represents the assumed 10% measurement uncertainty **B)** Frequency histograms (from 3.4e6 samples) comparing the probability distributions of the penalty parameters corresponding to the different network modules. All the distributions have deviated from the Laplace prior boundaries shown in pink, indicating that all the modules in the network were essential to fit this data.

**Table S1.**
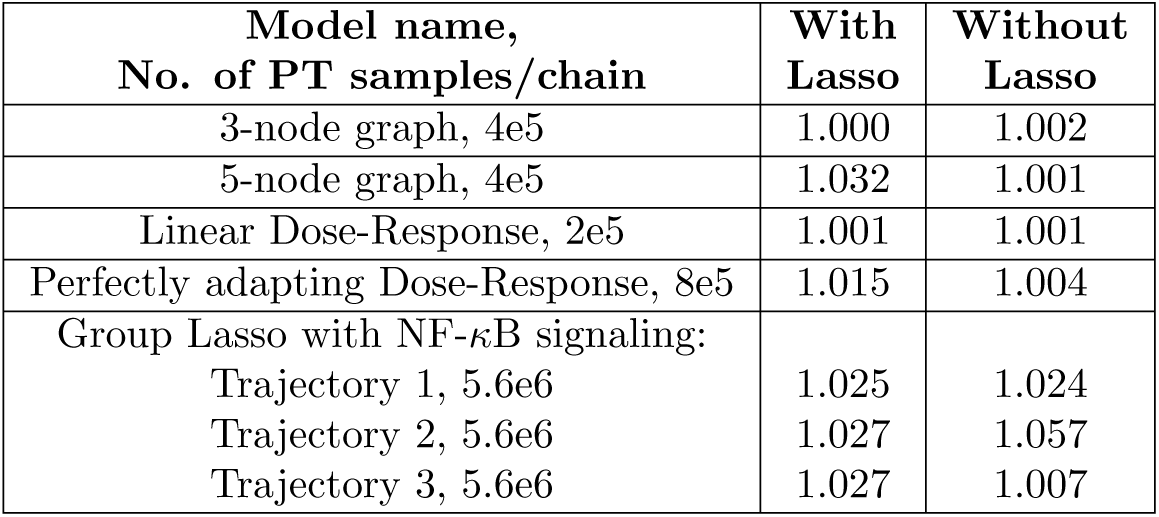
Maximum PSRF across all model parameters for each example shown up to 4 significant digits.

| Model name,<br>No. of PT samples/chain | With<br>Lasso | Without<br>Lasso |
| --- | --- | --- |
| 3-node graph, 4e5 | 1.000 | 1.002 |
| 5-node graph, 4e5 | 1.032 | 1.001 |
| Linear Dose-Response, 2e5 | 1.001 | 1.001 |
| Perfectly adapting Dose-Response, 8e5 | 1.015 | 1.004 |
| Group Lasso with NF-κB signaling: | | |
| Trajectory 1, 5.6e6 | 1.025 | 1.024 |
| Trajectory 2, 5.6e6 | 1.027 | 1.057 |
| Trajectory 3, 5.6e6 | 1.027 | 1.007 |

**Table S2.** MPSRF values for all PT runs in this study shown up to 4 significant digits.

| Model name,<br>No. of PT samples/chain | With<br>Lasso | Without<br>Lasso |
| --- | --- | --- |
| 3-node graph, 4e5 | 1.001 | 1.002 |
| 5-node graph, 4e5 | 1.031 | 1.010 |
| Linear Dose-Response, 2e5 | 1.003 | 1.004 |
| Perfectly adapting Dose-Response, 8e5 | 1.012 | 1.006 |
| Group Lasso with NF-κB signaling: | | |
| Trajectory 1, 5.6e6 | 1.074 | 1.094 |
| Trajectory 2, 5.6e6 | 1.120 | 1.200 |
| Trajectory 3, 5.6e6 | 1.129 | 1.014 |

**Table S3.** Step acceptance rates for the lowest temperature chain of all PT runs in this study.

| Model name,<br>No. of PT samples | With Lasso<br>(repeat 1, 2) | Without Lasso<br>(repeat 1, 2) |
| --- | --- | --- |
| 3-node graph, 4e5 | 0.24, 0.25 | 0.24, 0.25 |
| 5-node graph, 4e5 | 0.24, 0.23 | 0.24, 0.26 |
| Linear Dose-Response, 2e5 | 0.24, 0.22 | 0.24, 0.24 |
| Perfectly adapting Dose-Response, 8e5 | 0.24, 0.24 | 0.25, 0.25 |
| Group Lasso with NF-κB signaling, 5.8e6 | | |
| Trajectory 1 | 0.26, 0.26 | 0.23, 0.22 |
| Trajectory 2 | 0.26, 0.25 | 0.45, 0.26 |
| Trajectory 3 | 0.24, 0.25 | 0.26, 0.22 |

**Table S4.** Hard constraints in NF-*κ*B signaling fit

| Index | Constraint description | Constraint |
| --- | --- | --- |
| 1 | Exit rate of NF-κB -IκB from the nucleus is greater than free NF-κB | ke2a > ke1 |
| 2 | Exit rate of NF-κB -IκB from the nucleus is greater than free IκB | ke2a > ke2 |
| 3 | IKK mediated degradation of free cytosolic IκB is greater than the basal rate | kt1a > c4a |
| 4 | IKK mediated degradation of NF-κB bound cytosolic IκB is greater than the basal rate | kt2a > c5a |
| 5 | At equilibrium there is some NF-κB in the nucleus | $\frac{n\text{NF}\kappa\text{B}_{eq} + n\text{NF}\kappa\text{B} - \text{I}\kappa\text{B}_{eq}}{\text{TotalNF}\kappa\text{B}} \in [0.05, 0.5]$ |
| 6 | At least 1 TNFR molecule/cell is activated | $\min(\text{TNFRa}) > 1$ |
| 7 | At least 1 IKK molecule/cell is activated | $\min(\text{IKKa}) > 1$ |
| 8 | Equilibrium check before TNF stimulation | Rate of change for each model species < 1e-4 molecules/5e5 seconds |

**Table S5.** PSRF to show convergence of energy distributions when combining PT runs for NF-*κ*B signaling fit.

| Trajectory | Lasso group 1,<br>M=6, N=5.6e6 | Lasso group 2,<br>M=6, N=5.6e6 | Without Lasso<br>group 1, M=6,<br>N=5.6e6 | Without Lasso<br>group 2, M=6,<br>N=5.6e6 |
| --- | --- | --- | --- | --- |
| 1 | 1.041 | 1.016 | 1.049 | 1.021 |
| 2 | 1.139 | 1.134 | 1.011 | 1.003 |
| 3 | 1.028 | 1.009 | 1.001 | 1.003 |

**Table S6.** Swap acceptance rates for the two lowest temperature chains of all PT runs in this study.

| Model name,<br>No. of PT samples/chain | With Lasso<br>(repeat 1, 2) | Without Lasso<br>(repeat 1, 2) |
| --- | --- | --- |
| 3-node graph, 4e5 | 0.24, 0.33 | 0.25, 0.35 |
| 5-node graph, 4e5 | 0.33, 0.47 | 0.32, 0.26 |
| Linear Dose-Response, 2e5 | 0.26, 0.97 | 0.24, 0.24 |
| Perfectly adapting Dose-Response, 8e5 | 0.17, 0.37 | 0.25, 0.26 |
| Group Lasso with NF-κB signaling, 5.6e5 | | |
| Trajectory 1 | 0.28, 0.25 | 0.19, 0.25 |
| Trajectory 2 | 0.17, 0.17 | 0.29, 0.32 |
| Trajectory 3 | 0.16, 0.16 | 0.26, 0.28 |

## Acknowledgments

This work was funded by NIH grant R35-GM119462 to RECL, and by JRF via the NIGMS-funded (P41-GM103712) National Center for Multiscale Modeling of Biological Systems (MMBioS).

